# Evidence of genetic overlap between circadian preference and brain white matter microstructure

**DOI:** 10.1101/2020.11.09.374199

**Authors:** Luis M. García-Marín, Sarael Alcauter, Adrian I. Campos, Aoibhe Mulcahy, Pik-Fang Kho, Gabriel Cuéllar-Partida, Miguel E. Rentería

**Affiliations:** Genetic Epidemiology Lab, QIMR Berghofer Medical Research Institute, Brisbane QLD Australia; Instituto de Neurobiología, Universidad Nacional Autónoma de México, Querétaro, México; Faculty of Medicine, The University of Queensland, Brisbane QLD Australia; School of Biomedical Sciences, Faculty of Health, Queensland University of Technology, Brisbane, QLD, Australia

**Keywords:** Chronotype, circadian preference, white matter microstructure, diffusion tensor imaging, genetics, complex-traits, epidemiology, diffusion tensor imaging

## Abstract

**Study objective:** Previous neuroimaging studies have highlighted differences in white matter microstructure among individuals with different chronotypes, but it is unclear whether those differences are due to genetic or environmental factors.

**Methods:** Here we leverage summary statistics from recent large-scale genome-wide association studies (GWAS) of chronotype and diffusion tensor imaging (DTI) measures to examine the genetic overlap and infer causal relationships between these traits.

**Results:** We identified 29 significant pairwise genetic correlations, of which 13 also had evidence for a causal association. Negative genetic correlations were identified between chronotype and brain-wide mean, axial and radial diffusivities. When exploring individual tracts, ten negative genetic correlations were observed with mean diffusivities, 10 with axial diffusivities, 4 with radial diffusivities and 2 with mode of anisotropy. We found evidence for a possible causal association of chronotype with white matter microstructure in individual tracts including the posterior limb and retrolenticular part of the internal capsule; the genu and splenium of the corpus callosum and the posterior, superior and anterior regions of the corona radiata.

**Conclusions:** Our results suggest that *eveningness* is associated with variation in tract-specific white matter microstructure and, for an evening person, increases in axial and / or radial diffusivities may influence a higher mean diffusivity. These findings add to our understanding of circadian preference and its relationship with the brain, providing new perspectives on the genetic neurological underpinnings of chronotype’s role in health and disease.

**Statement of Significance:** Sleep is essential for a healthy brain function, particularly for neural organization and brain structure development. Individual chronotype differences have been associated with depression, schizophrenia, diabetes and obesity, among other conditions. Investigating the shared genetic aetiology between chronotype and white matter microstructure is essential to understand the neurological basis of individual variation in chronotype. In the present study, we show that tract-specific white matter microstructure is genetically correlated and causally associated with chronotype.

## Introduction

Circadian rhythms are fundamental physiological processes that occur daily in most living organisms. Humans are normally diurnal creatures, and human activity-rest patterns are endogenously controlled by biological clocks with a circadian (~24-hour) period. Notably, these cycles are known to have profound effects on fundamental molecular and behavioural processes, including the regulation of hormone levels, body temperature and sleep-wake patterns.[1]

The expression of circadian preference, often referred to as chronotype, is the natural phenomena accounting for the variation in an individual’s preference for early or late sleep and activity driven by sleep-wake cycles due to internal circadian rhythms. Furthermore, chronotype defines whether an individual is a *morning person*, implying a preference for going to bed and waking up early, or an *evening person*, meaning someone who prefers later sleep and waking up times. Normal variation in chronotype encompasses sleep-wake cycles that are two to three hours later in evening types than morning types. Extremes outside of this range can result in difficulty in participating in regular work, school, and social activities and are thus considered to have a sleep disorder.[2] A considerable proportion of variance in chronotype is due to factors such as age, gender and the exposure to different levels of light. However, genetic variation also accounts for chronotype differences and twin studies have estimated its heritability at ~50%. [3–6]

Adequate sleep is vital for healthy brain function and physiological systems.[7] The *restoration theory* states that sleep is essential to replete and restore the cellular components necessary for biological functions.[8] In contrast, the *brain plasticity theory* describes the vital neural reorganisation and development of the brain’s diverse structures and functions that occurs during the restorative state.[8]

Circadian disorders have been linked to several health outcomes in humans, and there is a growing interest in the implications of chronotype on health. For instance, in genetic studies, being a “lark”, or *morning person* has been negatively correlated on the genetic level with depression and schizophrenia,[2] and positively correlated with subjective well-being.[2] Also, observational studies show associations between being an evening person and risk for diabetes,[9] metabolic syndrome,[9] sarcopenia[9] and obesity.[10] Further, Mendelian randomisation analyses indicate that being a morning person may lead to better mental health.[2]

Recent studies have shown that brain structure as determined by neuroimaging methods is highly heritable,[11,12] and the growing use of imaging techniques in genetic studies has opened up new lines of investigation for several phenotypes. For instance, neuroimaging techniques such as computerized tomography (CT), magnetic resonance imaging (MRI) and positron emission tomography (PET) have proven useful to gain a better understanding of insomnia,[13] schizophrenia,[14,15] depression[14,16] and Alzheimer’s disease[14], among other conditions.

Diffusion-weighted imaging is a non-invasive neuroimaging technique that can be used to characterise the brain’s white matter integrity in acute sleep deprivation, insomnia and chronotype.[17,18] In particular, diffusion tensor imaging (DTI) refers to the implementation of a tensor model to explore the main directions (eigenvectors of the model) and magnitude (eigenvalues) of water diffusivity[19], which are sensitive to the underlying microstructure.[20] Water diffusivity parameters are combined into several indices to facilitate their interpretation. For instance, axial diffusivity (AD) is the first eigenvalue and is related to the diffusivity along the main direction of water diffusion[21]; *radial diffusivity* (RD), is the average of the second and third eigenvalues. Thus, it represents the diffusivity in the secondary directions of water diffusion [21]; *mean diffusivity* (MD) is the average of the three eigenvectors, representing the average water diffusivity[21]. Further, *fractional anisotropy* (FA) is the normalised variance of the three eigenvalues. Therefore, it is increased if there is a main direction of diffusivity, and it is close to zero if water diffusivity is similar for any direction (isotropic)[21]. The *mode of anisotropy* (MO) characterises the general architecture of the tensor, from being disc-like, such as in brain regions with fibres crossing, to being highly tubular, as in regions with unidirectionally oriented fibres[22].

DTI, being a useful tool to explore brain regions with single fibre orientation, is used to infer histological changes in both normal development and pathology. For instance, AD is associated with axonal integrity, while RD is related to axonal density, myelin integrity, axonal diameter, and fibre coherence.[20] DTI has been used to characterise white matter integrity in sleep disorders including insomnia[23]^,[24]^ and narcolepsy,[25,26] in both cases showing decreased FA in the prefrontal regions and the anterior internal capsule inducing hypoactivation of the prefrontal cortex, which in turn is associated with daytime fatigue and comorbid psychiatric and cognitive deficits [27].

Genome-wide association studies (GWAS) of neuroimaging measures have uncovered 213 independent significant genetic variants associated with 90 DTI parameters.[28] Also, recent studies have described associations between sleep quality and the white matter integrity of several brain regions, including the cingulum, corpus callosum and frontal association tracts, further revealing the intrinsic relation between sleep and brain structural and functional integrity.[29–31]

Moreover, neuroimaging studies have identified white matter differences in *evening people*, especially in the frontal and temporal lobes, cingulate gyrus and corpus callosum.[32] Studies investigating the disruption of circadian cycles show that a day of wakefulness is associated with increases in white matter fractional anisotropy due to radial diffusivity reductions.[33,34] In contrast, sleep deprivation is associated with a lessening of fractional anisotropy driven by reductions in axial diffusivity.[33] Thus, suggesting that wakefulness-related effects may regulate white matter.[34]

It is unclear whether the association between DTI and sleep is due to genetic factors, and if so, whether pleiotropic or causal effects drive the association. In the present study, we address this question by leveraging the availability of GWAS summary statistics datasets to estimate whole-genome genetic correlations and use Mendelian randomisation analysis to assess causality between DTI measures and circadian preference. We demonstrate that an evening chronotype is genetically associated with variation in 29 white matter microstructure parameters including increases in mean, axial and radial diffusivities and mode of anisotropy among specific tracts in the internal capsule, the corona radiata and the corpus callosum. MR analyses provide evidence for 13 causal relationships between chronotype and DTI parameters. These results provide new insights into the relationship between chronotype and white matter structure and contribute to our understanding of the genetic and neurological factors underlying chronotype.

## Methods

### Discovery GWAS datasets for DTI brain measures

The present study used summary statistics from genome-wide association studies with European ancestry for white matter microstructure, particularly diffusion tensor imaging, objectively measured with magnetic resonance imaging (MRI). Summary statistics include allele frequency, effect size, standard error, as well as the p-value of every genetic variant that was tested on the trait of interest. Frequently, published GWAS studies have made their summary statistics available to the scientific community to provide access to genetic data, which in turn has increased the number of available resources for researchers to advance our understanding of genetic factors underlying a wide range of phenotypes. The GWAS summary statistics used here were obtained from samples with European ancestry (*N = 17,706*) from a public repository as reported in its corresponding publication (Zhao et al. 2019).[28] White matter microstructure parameters included mean diffusivities (MD), axial diffusivities (AD), radial diffusivities (RD) mode of anisotropies (MO) and fractional anisotropy (FA). GWAS were adjusted for age, age^2^, sex, age*sex interaction, age^2^*sex interaction, and the top 10 genetic principal components.[28]

### Discovery GWAS dataset for circadian preference

This study used summary statistics from genome-wide association studies for self-reported chronotype (*N = 449,734*). These GWAS summary statistics were obtained from samples with European ancestry from the repository of its corresponding publication (Jones et al. 2019).[2] Chronotype GWAS summary statistics were adjusted for age, sex, study centre and genotyping array.[2] No age cut-offs were applied.[2]

### Genetic correlations analyses

We performed LD score regression[35] using GWAS summary statistics as implemented in the Complex Trait Genomics Virtual Lab (CTG-VL, http://genoma.io)[36] to estimate genetic correlations between chronotype and white matter microstructures. Benjamini-Hochberg’s False Discovery Rate (FDR < 5%) was used to account for multiple testing.

### Mendelian randomisation

Mendelian randomisation (MR) is a method used in epidemiology to assess causal relationships between phenotypes, environmental exposures or disease outcomes.[37–39] We used two-sample Generalised Summary-data-based Mendelian Randomisation (GSMR),[39] a method that uses genetic instruments strongly associated (i.e genome-wide significant SNPs) with the outcome, to assess a possible causal relationship between chronotype and 29 DTI parameters which displayed a significant genetic correlation. P-values were corrected for multiple testing using Benjamini-Hochberg’s False Discovery Rate (FDR < 5%). Both corrected and uncorrected p-values are reported.

## Results

### Genetic correlations

From the 110 DTI parameters that were analyzed, we identified 29 significant negative genetic correlations (FDR < 5%) between chronotype and DTI parameters (**Table 1**). Negative correlations with chronotype imply positive correlations with eveningness, that is, a preference for late sleep patterns and activity. The average MD, AD, and RD for all tracts showed significant negative genetic correlations with chronotype. When exploring the individual tracts (**Figure 1**), 10 out of 26 genetic correlations involved mean diffusivity (MD), 10 included axial diffusivity (AD), four implicated radial diffusivity (RD) and two involved mode of anisotropy (MO). No significant genetic correlations were identified with fractional anisotropy (FA).

**Table 1.**
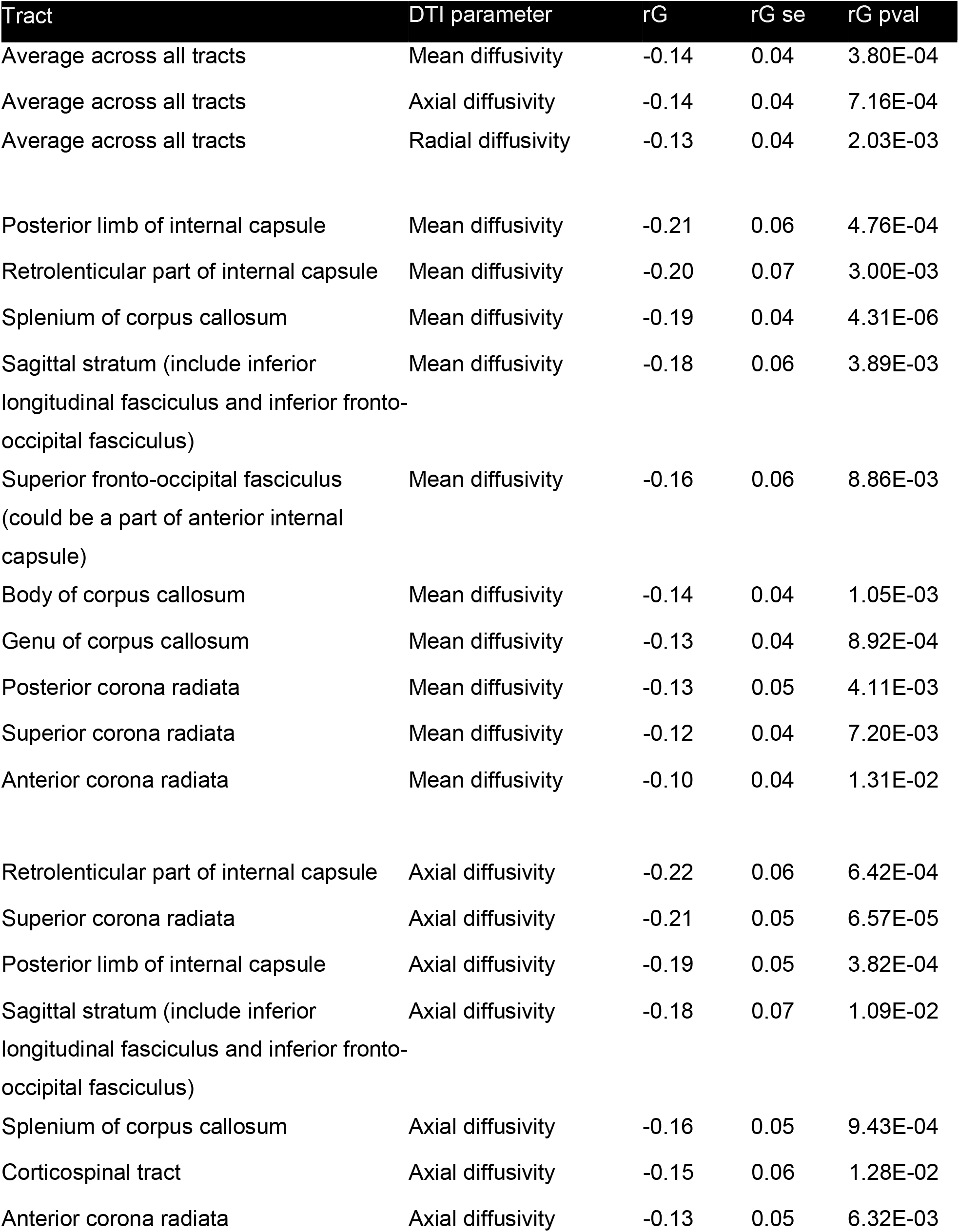
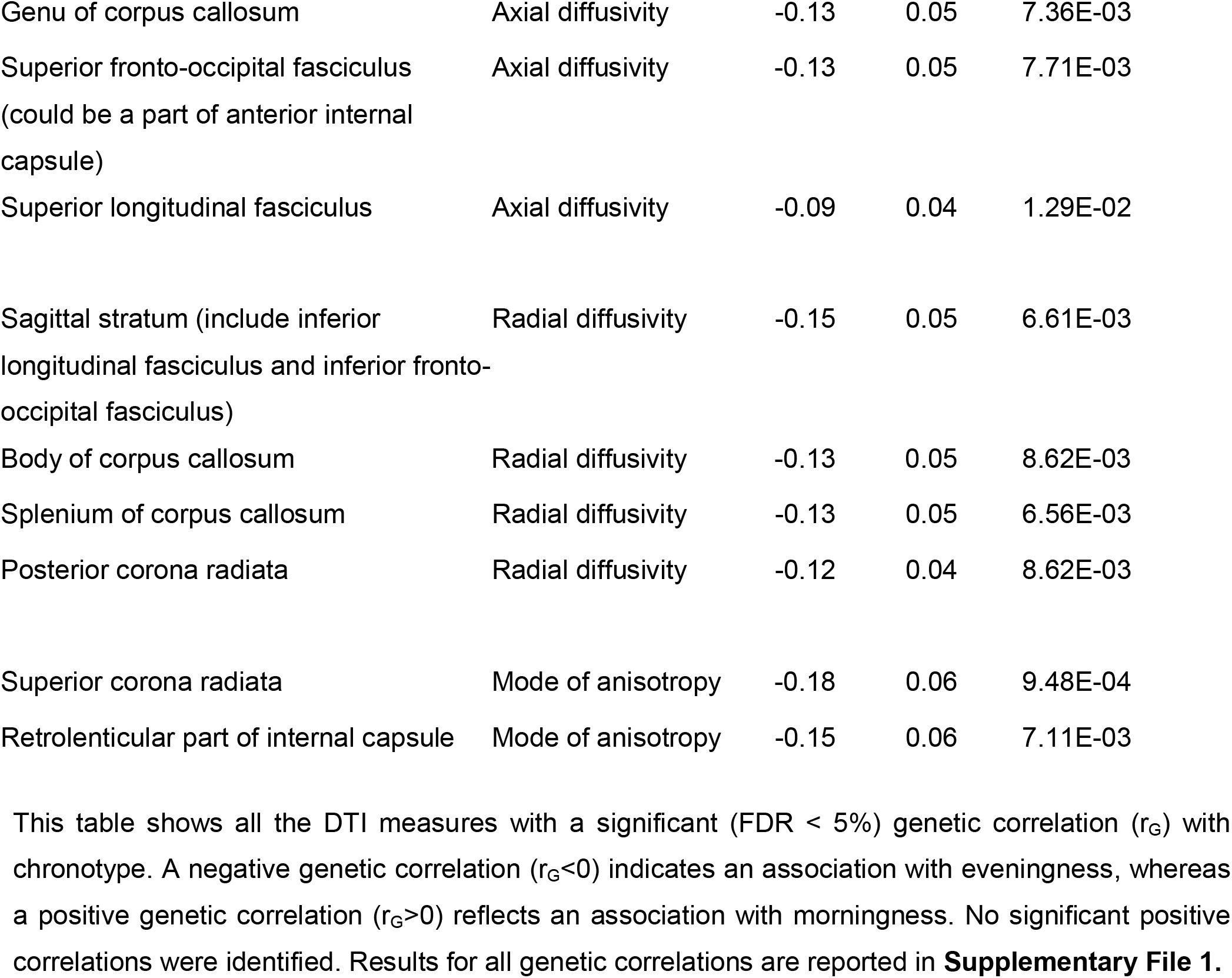
White matter microstructure measures genetically correlated with chronotype.

**Figure 1.**
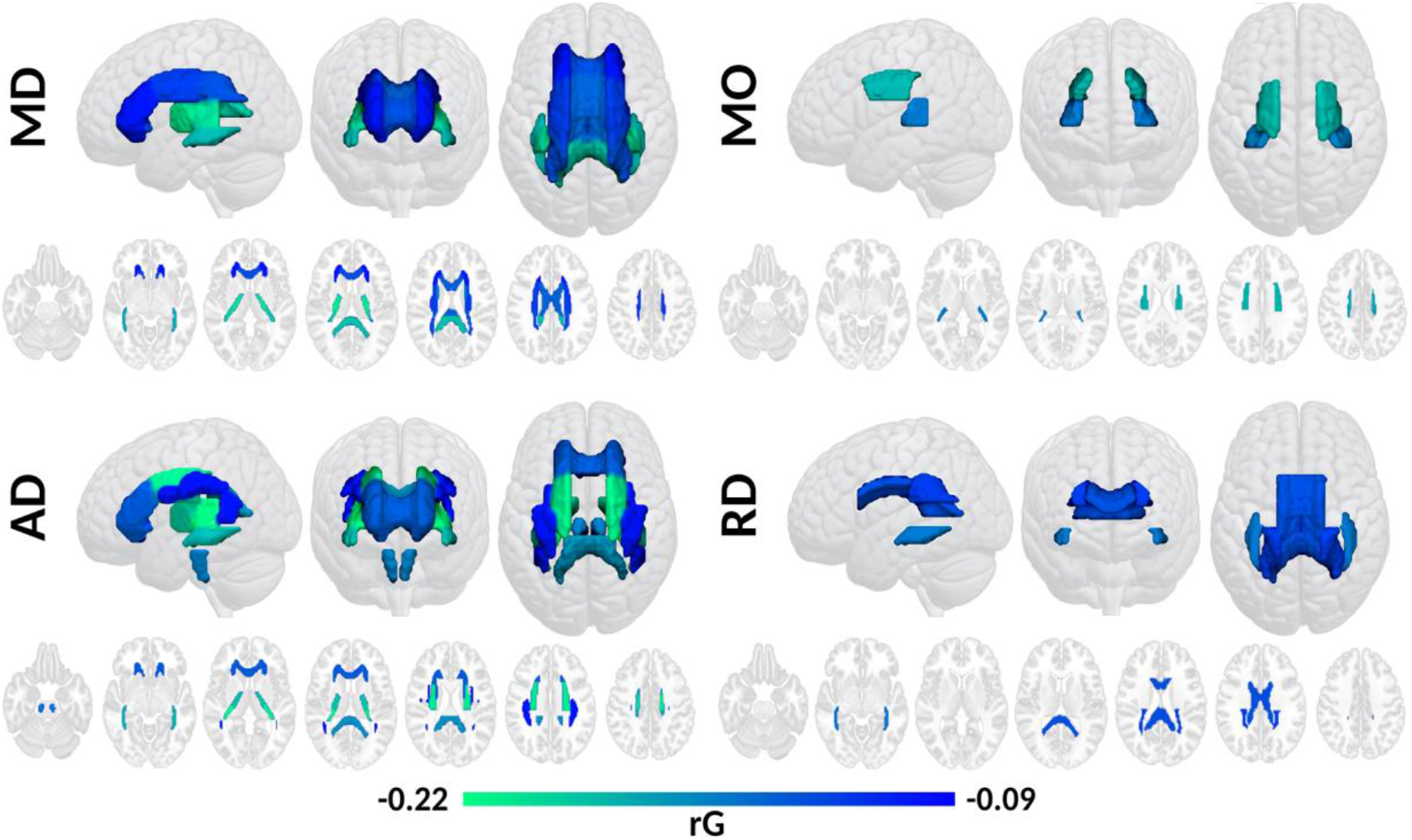
Significant genetic correlations between chronotype and DTI parameters classified by individual tracts including (MD) mean diffusivity, (MO) mode of anisotropy, (AD) axial diffusivity and (RD) radial diffusivity. All genetic correlations are negative. Negative correlations with chronotype imply positive correlations with eveningness.

For MD, the strongest genetic correlations with chronotype were observed for the internal capsule, particularly in its posterior limb and the retrolenticular region. Further, the most significant genetic correlation was identified in the splenium of the corpus callosum, but the genu and body of the corpus callosum were also genetically correlated with being an evening person. In addition, the posterior, superior and anterior regions of the corona radiata were also genetically correlated with chronotype. A similar pattern was observed for the sagittal stratum and the superior fronto-occipital fasciculus.

Regarding AD, robust genetic correlations were identified in the corona radiata with the most significant one in the superior region. The anterior region of the corona radiata was also genetically correlated with chronotype. Moreover, genetic correlates were observed for the retrolenticular regions and the posterior limb of the internal capsule, followed by the splenium and genu of the corpus callosum. More genetic correlations were uncovered for the superior fronto-occipital fasciculus, the superior longitudinal fasciculus, the sagittal stratum and the corticospinal tract.

Genetic correlations identified between chronotype and RD included the splenium and body of the corpus callosum, the posterior region of the corona radiata and the sagittal stratum. In contrast, MO displayed moderate genetic correlates with chronotype in the superior region of the corona radiata and the retrolenticular section of the internal capsule.

### Mendelian randomisation

To further explore the associations between chronotype and DTI, we performed a two-sample Generalised Summary-data-based Mendelian Randomisation analysis (GSMR)[39] and assessed the potential of the 29 genetic correlations we identified to be explained by a causal association (**Supplementary File 2**).

We observed 13 one-way causal relationships in which an evening chronotype influences increases in mean, axial and radial diffusivities (**Table 2**). GSMR results suggest that eveningness exerts a causal effect on higher MD for the superior fronto-occipital fasciculus and the reticular region and posterior limb of the internal capsule. Further, increases in mean and axial diffusivities for the posterior corona radiata and the splenium of the corpus callosum were found to be causally influenced by an evening chronotype. A similar pattern was observed for mean and radial diffusivities for the superior and anterior regions of the corona radiata. Also, a one-way causal relationship was identified for eveningness increasing RD for the superior longitudinal fasciculus.

**Table 2.**
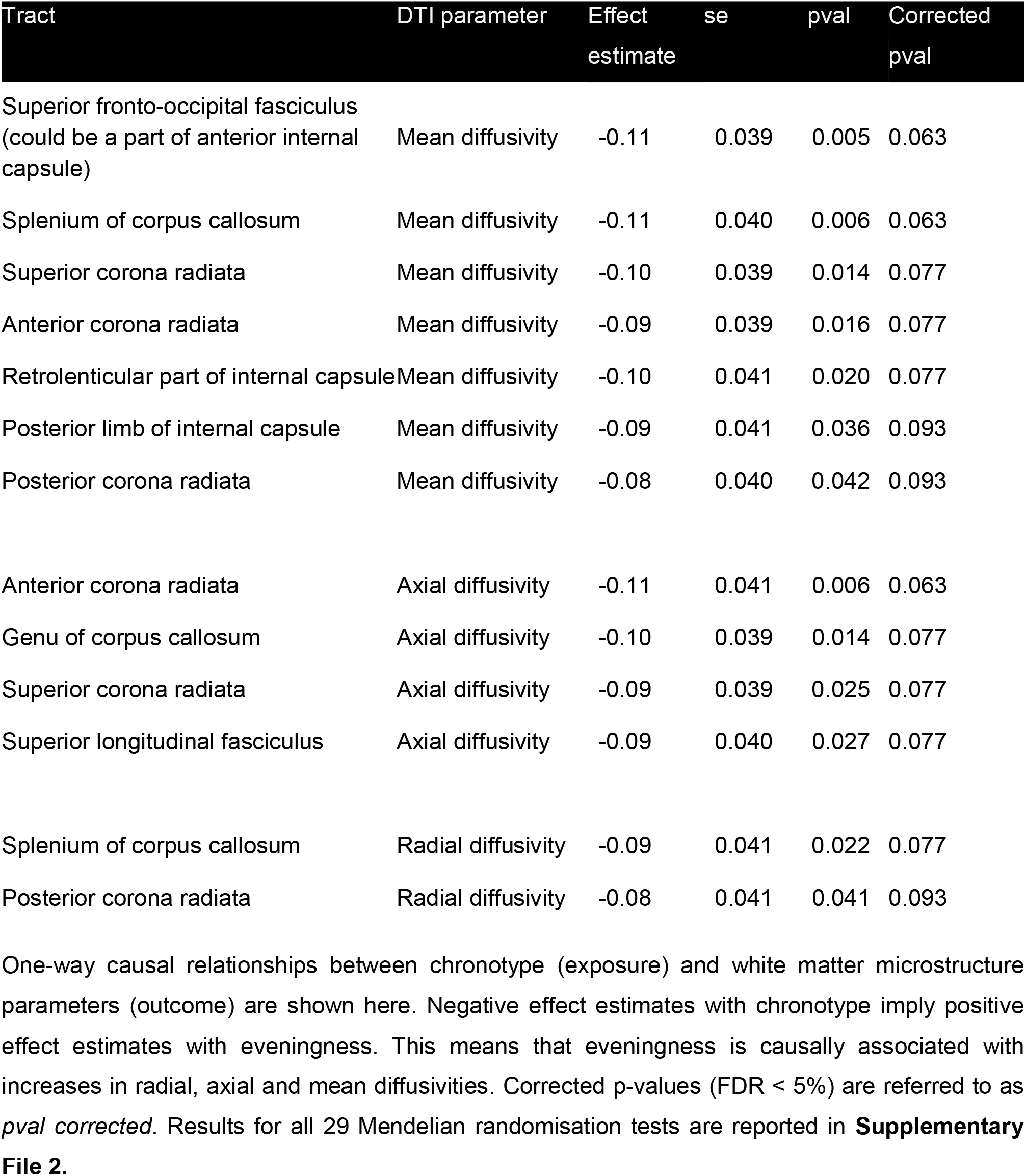
Mendelian randomization results for chronotype and DTI.

| Tract | DTI parameter | Effect estimate | se | pval | Corrected pval |
| --- | --- | --- | --- | --- | --- |
| Superior fronto-occipital fasciculus (could be a part of anterior internal capsule) | Mean diffusivity | -0.11 | 0.039 | 0.005 | 0.063 |
| Splenium of corpus callosum | Mean diffusivity | -0.11 | 0.040 | 0.006 | 0.063 |
| Superior corona radiata | Mean diffusivity | -0.10 | 0.039 | 0.014 | 0.077 |
| Anterior corona radiata | Mean diffusivity | -0.09 | 0.039 | 0.016 | 0.077 |
| Retrolenticular part of internal capsule | Mean diffusivity | -0.10 | 0.041 | 0.020 | 0.077 |
| Posterior limb of internal capsule | Mean diffusivity | -0.09 | 0.041 | 0.036 | 0.093 |
| Posterior corona radiata | Mean diffusivity | -0.08 | 0.040 | 0.042 | 0.093 |
| Anterior corona radiata | Axial diffusivity | -0.11 | 0.041 | 0.006 | 0.063 |
| Genu of corpus callosum | Axial diffusivity | -0.10 | 0.039 | 0.014 | 0.077 |
| Superior corona radiata | Axial diffusivity | -0.09 | 0.039 | 0.025 | 0.077 |
| Superior longitudinal fasciculus | Axial diffusivity | -0.09 | 0.040 | 0.027 | 0.077 |
| Splenium of corpus callosum | Radial diffusivity | -0.09 | 0.041 | 0.022 | 0.077 |
| Posterior corona radiata | Radial diffusivity | -0.08 | 0.041 | 0.041 | 0.093 |

One-way causal relationships between chronotype (exposure) and white matter microstructure parameters (outcome) are shown here. Negative effect estimates with chronotype imply positive effect estimates with eveningness. This means that eveningness is causally associated with increases in radial, axial and mean diffusivities. Corrected p-values (FDR < 5%) are referred to as *pval corrected*. Results for all 29 Mendelian randomisation tests are reported in **Supplementary File 2**.

## Discussion

This study provides insights into the relation between chronotype and DTI. We identified 29 significant negative genetic correlations between chronotype and DTI parameters, including MD, AD, RD, and MO. Of those, 13 were found to be indicative of a causal relationship. These results show that changes in white matter microstructural organisation associated with an evening chronotype are explained by variation in MD, AD, RD and MO tracts.

Recently, there has been a growing interest in the genetic architecture of chronotype and its role in health and disease.[2,40] For instance, being a morning person has been proven to be genetically correlated with metabolic traits such as body mass index (BMI)[2], type 2 diabetes[2] and fasting insulin.[2] Further, phenotypes involving depression such as *ever had prolonged feelings of sadness or depression [40]* and *ever depressed for a whole week* [40] have shown negative genetic correlations with morningness. In contrast, diseases of the musculoskeletal system and connective tissue have shown a positive genetic correlation with being a morning person.[40]

In the present study, no positive genetic correlations were identified. Nonetheless, MD, AD, RD and MO showed significant negative genetic correlations with chronotype. Specifically, negative correlations with chronotype imply that DTI parameters are positively correlated with eveningness. That is, the eveningness chronotype is associated with higher average MD, AD, and RD for all tracts, but with tract-specific distributions for each DTI parameter. Our results are highly consistent with previous findings[32] describing white matter microstructural differences in several brain regions including the corpus callosum, corona radiata, internal capsule, frontal and temporal lobes when comparing evening to morning and intermediate chronotypes.

We identified genetic correlations between an evening chronotype and MD in the entire corpus callosum and corona radiata, as with the posterior segments of the internal capsule. Increased MD may be associated with decreased tissue density, altered properties of the myelin, axonal and neuronal membrane, as well as altered tissue organization and shape of glia and neurons.[20,41] Further, in our study, the posterior region of the corona radiata also showed increased RD, but no associations were detected with AD. An increase in AD, but not in RD, was identified in the posterior portion of the internal capsule for the evening chronotype. Similarly, the anterior part of the corona radiata showed genetic correlations between evening chronotype and AD, but not with RD. AD is usually associated with axonal membranes or axonal density,[42] while RD is typically associated with myelin density.[42] However, both measures could be affected by multiple tissue factors, including the presence of glial tissue, edema or multiple crossing fibre tracts.[43] Although there is not a straightforward interpretation of the underlying tissue characteristics based on DTI parameters, our results suggest an association between white matter microstructure and chronotype and we speculate that increments in AD, RD, or both, could influence higher MD in the evening chronotype. However, more research is needed to disentangle the intricate relationship between AD, RD and MD.

Higher MO is associated with a more tubular shape of the diffusion tensor, consistent with the presence of a predominant fibre population. Our results showed that higher MO was associated with a genetic predisposition to the *evening chronotype* in regions with expected crossing fibres (superior corona radiata and the retrolenticular portion of the internal capsule).

Such increase in MO may be attributed to decreased quality (less densely packed fibres and axonal density) of a portion of the crossing fibres, resulting in a predominant fibre population, also consistent with the increase of AD and MD in the same regions.[44,45]

Evening chronotype has been associated with lower sleep quality,[46,47] which in turn, has been related to increased MD in the hippocampus, basal ganglia, anterior corpus callosum and prefrontal white matter.[29,30] Our findings show similar patterns of white matter integrity in evening chronotype and further contribute to identifying an underlying genetic component between chronotype and MD regions.

Limitations of the present study must be recognised. Our analyses only used data from participants with European ancestry from the UK Biobank cohort. Since previous studies have pointed out racial differences in circadian rhythms,[48,49] the generalizability of our results may be limited and should be addressed with caution. Furthermore, Mendelian randomisation methods rely heavily on the statistical power of the samples, which in turn is tied to the availability of sufficient genetic instruments (i.e. genome-wide significant loci).[40,50] In the present study, individual DTI parameters showed scarce genome-wide significant loci, lessening the statistical power of GSMR. Nonetheless, our findings still provide evidence for causal associations suggesting chronotype is causal for white matter microstructure variation and thus, uncover meaningful insights and hypotheses for future studies to confirm when more powerful genetic statistical instruments and techniques become available.

In summary, we provide evidence for 29 genetic correlations between chronotype and brain white matter structure, of which 13 were found to be indicative of a causal relationship. Our results confirmed findings from previous studies and uncovered insights into the relationship between DTI parameters and chronotype. For instance, we reveal that increments in MD in the entire corona radiata and corpus callosum are associated with an evening chronotype. Further, we identified associations between evening chronotype and increments in the anterior and posterior parts of the corona radiata in RD and AD, respectively. Also, we found the retrolenticular part of the internal capsule to have higher MD, AD and MO among individuals with evening chronotype. Altogether, our results contribute to a better understanding of the relationship between the brain and circadian preference. Future studies should aim at disentangling the relationships between individual tracts to advance our understanding of the human brain and its role in sleep, activity and resting patterns.

## Ethics statement

This study was approved by the Human Research Ethics Committee of the QIMR Berghofer Medical Research Institute.

## Supplementary Files

**Supplementary File 1.** Genetic correlations between white matter microstructure measures and chronotype.

**Supplementary File 2.** Mendelian randomisation results for white matter microstructure measures and chronotype.

